## supplementary information for "The reciprocal regulation between mitochondrial-associated membranes and Notch signaling in skeletal muscle atrophy"

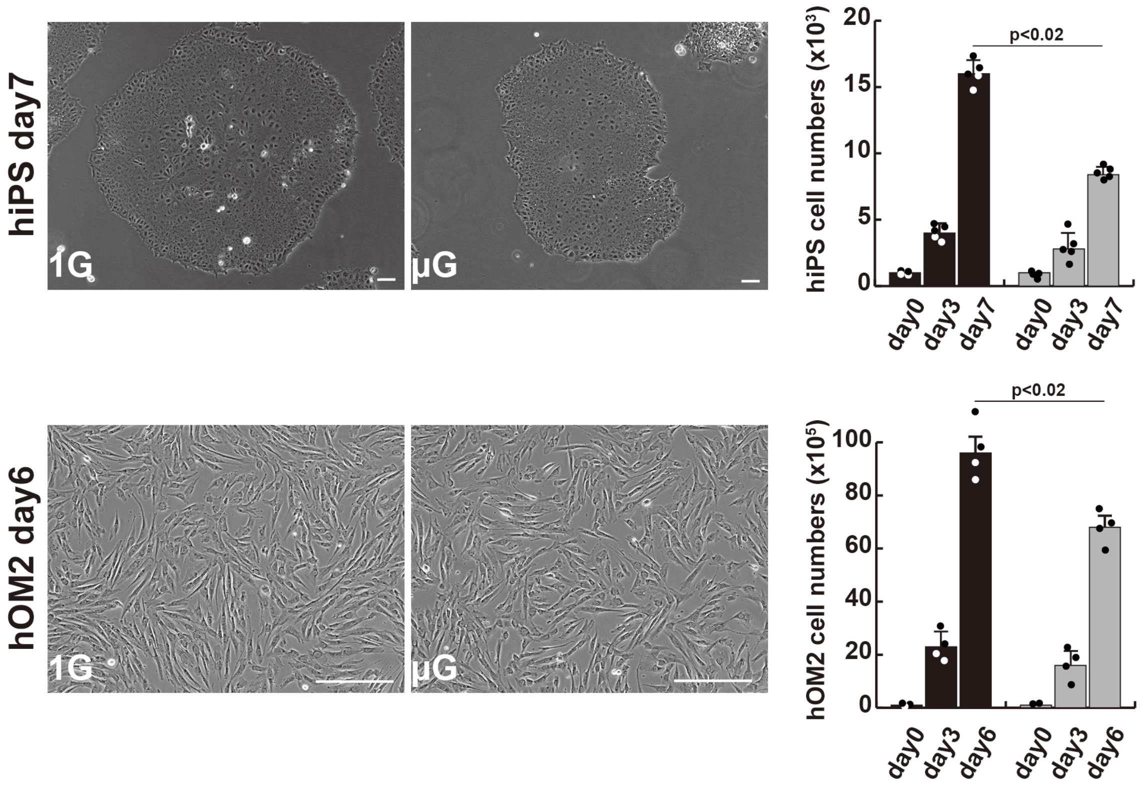


**Supplementary file 1. The effects of microgravity on cell proliferation in human iPS cells and hOM2 primary myogenic cells.** Phase contrast images showing cell morphology diagram after approximately 1 week of cell culture in normal (1G) or microgravity environment (μG) (left panels), and histograms showing the number of proliferating cells in cell culture (right). Scale bars; 100 μm. All error bars indicate ±SEM (n=5). *P*-values are determined by one-way ANOVA and Tukey’s test for comparisons.

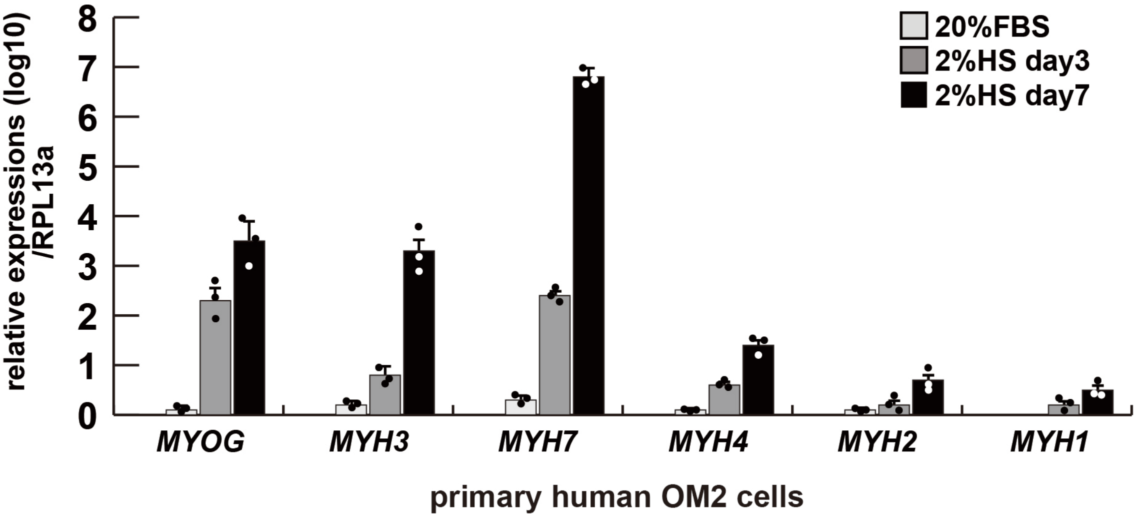


**Supplementary file 2. Comparative expression levels of myogenic genes in differentiated human primary hOM2 myogenic cells.** Differentiated primary human OM2 cells were examined by RT-qPCR for transcription of the myogenic differentiation markers MYOGENIN (*MYOG*), early differentiated Myosin Heavy Chain 3 (*MYH3*), and the type 1 slow muscle marker *MYH7*, as well as the fast muscle markers *MYH4*, *MYH2*, and *MYH1* (type 2b, type 2a, and type 2X, respectively), expressed relative to transcripts for the ribosomal protein RPL13a. FBS; fetal bovine serum, HS; horse serum

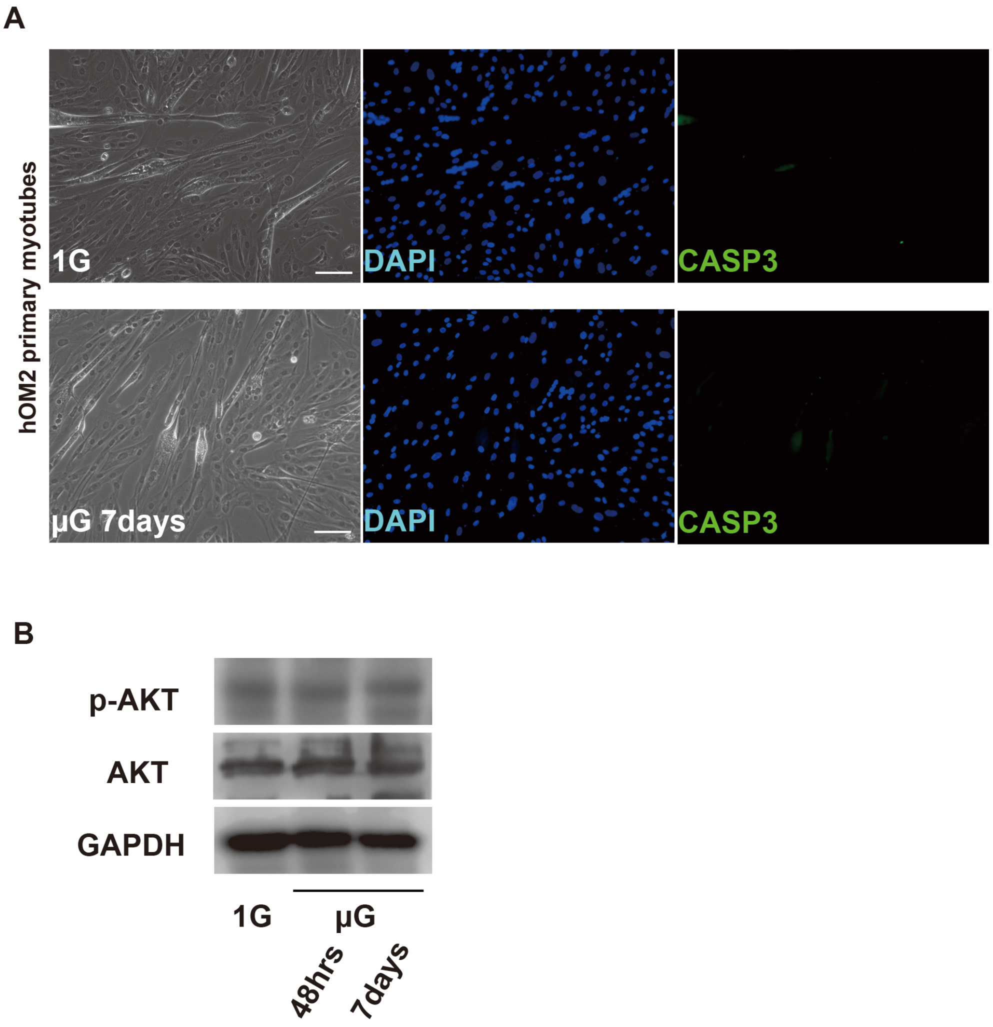


**Supplementary file 3. Apoptotic assays in differentiated hOM2 myotubes under microgravity.** (A) Phase contrast (left panels) and Immunofluorescent (middle and right panels) images of Caspase-3 (CASP3, green) and DAPI (blue) staining with differentiated hOM2 cells on normal (1G) or microgravity conditions (µG for 7 days). Scale bar; 100 µm. (B) Western Blotting with differentiated hOM2 cells against anti-phospho-AKT (p-AKT) antibody with or without the treatment of microgravity. GAPDH (whole cell) and AKT (pan) are used as loading controls.

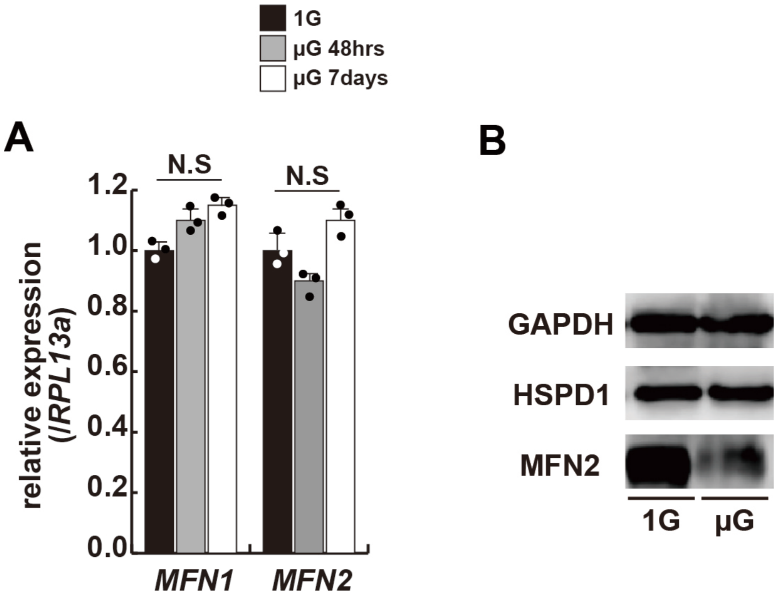


**Supplementary file 4. Transcription and translational levels of Mfn1/2 expression in differentiated hOM2 myotubes under microgravity.** (A) Relative transcript levels of *MFN1* and *MFN2* in differentiated hOM2 cells on normal (1G) or microgravity conditions (μG 48hrs and 7 days). (B) Western Blotting with differentiated hOM2 cells against anti-MFN2 antibody with or without the treatment of microgravity. GAPDH (whole cell) and HSPD1 (nuclear) are used as loading controls.

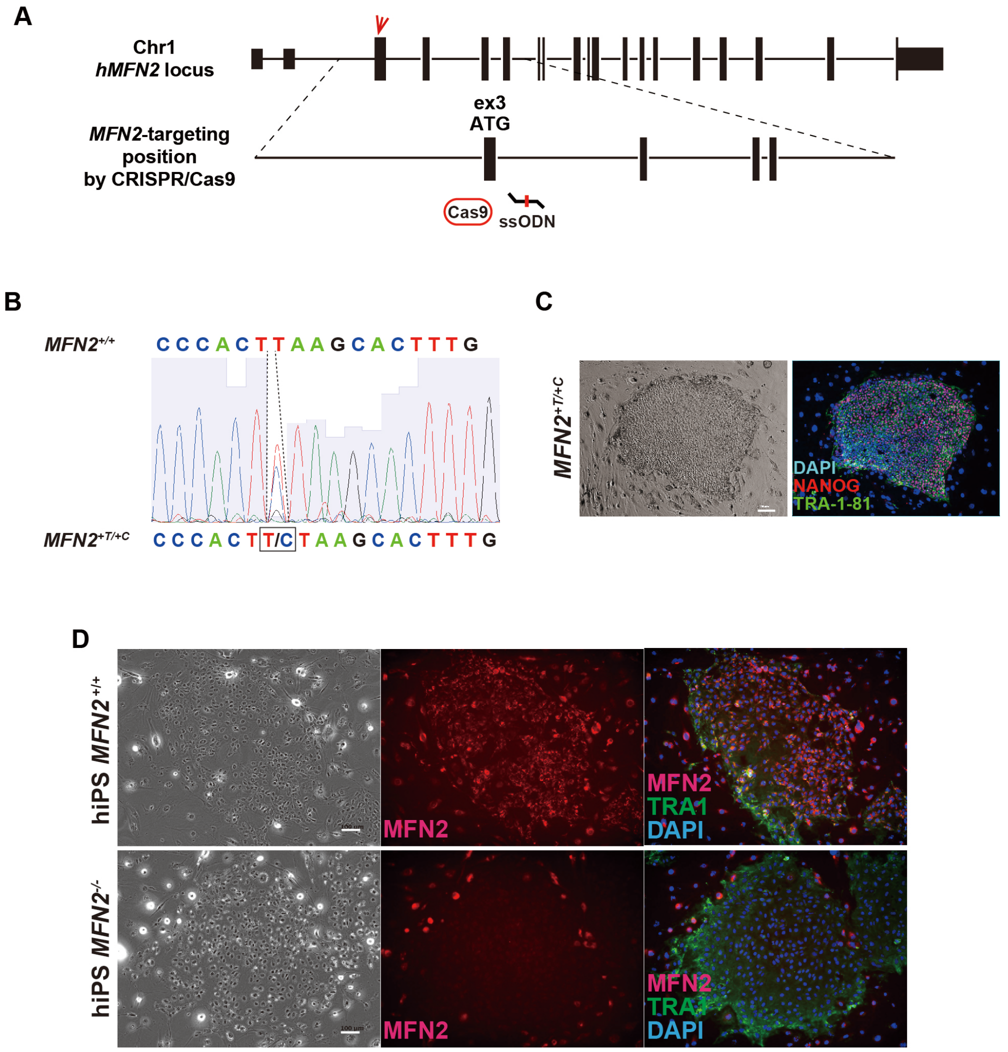


**Supplementary file 5. The generation of hMFN2-deficient human iPS cells**. (A) a schematic diagram of chromosome1 where the human MFN2 gene resides, and the target site of MFN2 exon 3 containing start ATG sequences treated by the CRISPR/Cas9 system with single-strand oligodeoxynucleotide (ssODN), which have knock-in sequences (additional 1 base T or C). (B) The result of DNA sequencing showing single nucleotide addition (T/ or C, lower sequence) sequencing using knock-in human iPS cells treated with CRISPR/Cas9 and ssODN. (C) Immunofluorescent analysis with *MFN2* knock-in human iPS cells (*MFN2*$^{+T/+C}$) to check the undifferentiated state. NANOG (red), TRA-1-81 (TRA1, Green), and DAPI (blue). Scale bar; 100 µm. (D) Immunofluorescent analysis with wildtype (*MFN2*$^{+/+}$) and *MFN2*-knockin human iPS cells (*MFN2*$^{-/-}$ as *MFN2*$^{+T/+C}$) to check the expression of MFN2 protein. MFN2 (red), TRA1 (green), and DAPI (blue). Scale bar; 50 µm.

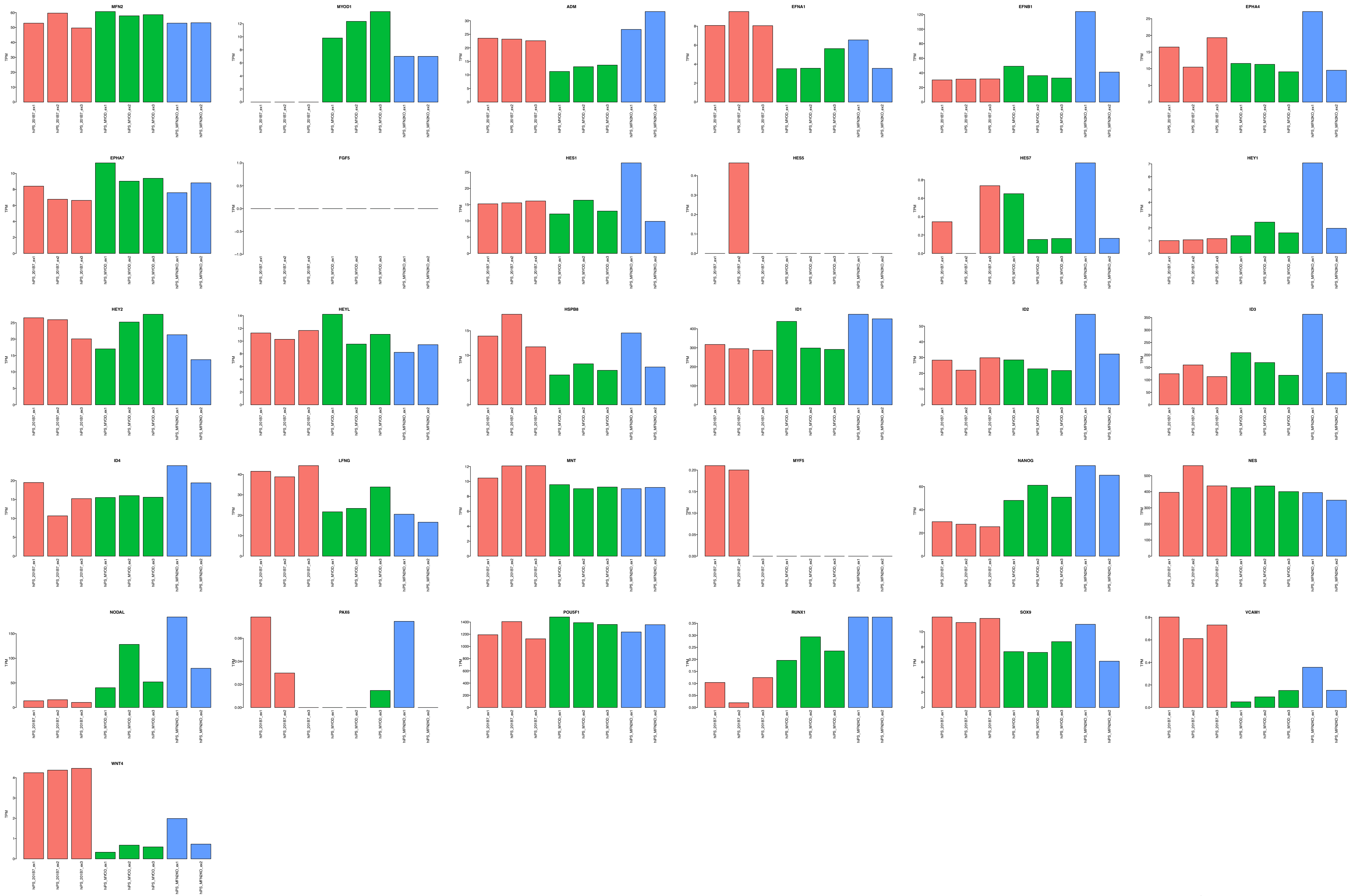


**Supplementary file 6. NGS analyses comparing normal hiPS (red, green) to *MFN2*$^{-/-}$ hiPS (blue) cells**

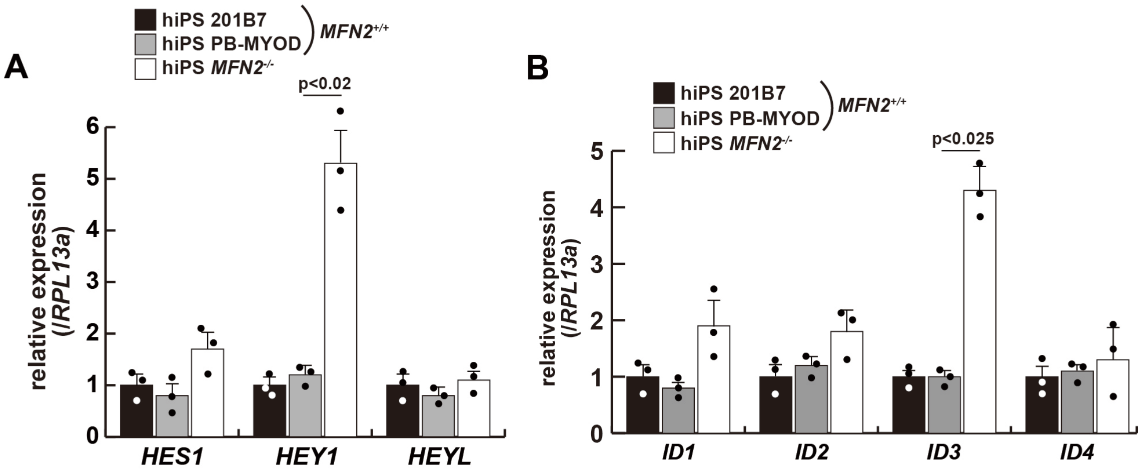


**Supplementary file 7. The relative expressions of Notch-related genes in MFN2-deficient human iPS cells.** Transcripts of the HES family (A) and ID family (B), which are downstream of the Notch signaling pathway, were evaluated in wildtype (201B7 or PB-MYOD) and MFN2-deficient (*MFN2*$^{-/-}$) human iPS cells. All error bars indicate ±SEM (n=3). *P*-values are determined by one-way ANOVA and Tukey's test for comparisons.

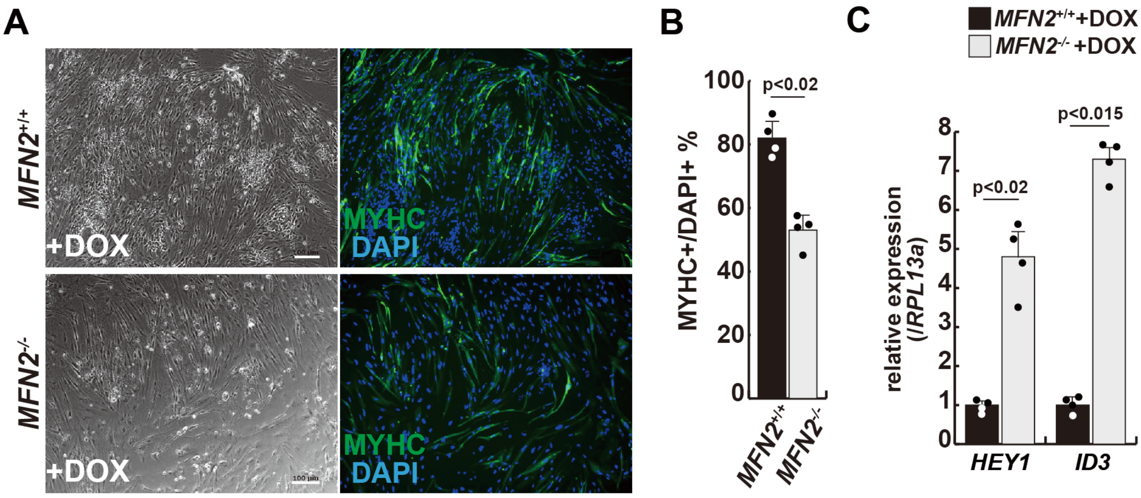


**Supplementary file 8. Differentiation of myogenic cells derived from MFN2-deficient human iPS cells.** (A) Phase contrast (left panels) and Immunofluorescent (right panels) images of Myosin Heavy Chain (MYHC, green) and DAPI (blue) staining with control and mutant (*MFN2*$^{-/-}$) differentiated human iPS cells after the administration of Doxycycline (DOX). (B) The percentage of MYHC/DAPI positive cells per fixed area in differentiated cells derived from wildtype or MFN2-deficient human iPS cells. (C) Relative level of *HEY1* and *ID3* transcripts in induced cells derived from human iPS cells with DOX treatment (*MFN2*$^{+/+}$ or *MFN2*$^{-/-}$ +DOX). All error bars indicate ±SEM (n=4). *P*-values are determined by non-parametric Wilcoxon tests for comparisons.

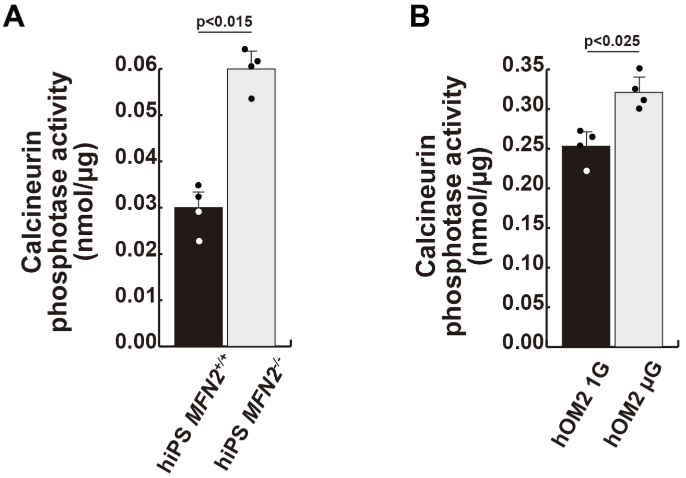


**Supplementary file 9. Calcineurin activity in human iPS cells and differentiated muscle cell cultures.** (A) Calcineurin activity was monitored in wildtype (*MFN*$^{+/+}$) and MFN2-deficient (*MFN2*$^{-/-}$) human iPS cells. (B) Differentiated human hOM2 cells were utilized in both normal gravity (1G) and microgravity (μG) conditions. All error bars indicate ±SEM (n=4). *P*-values are determined by non-parametric Wilcoxon tests for comparisons. *P<0.05.

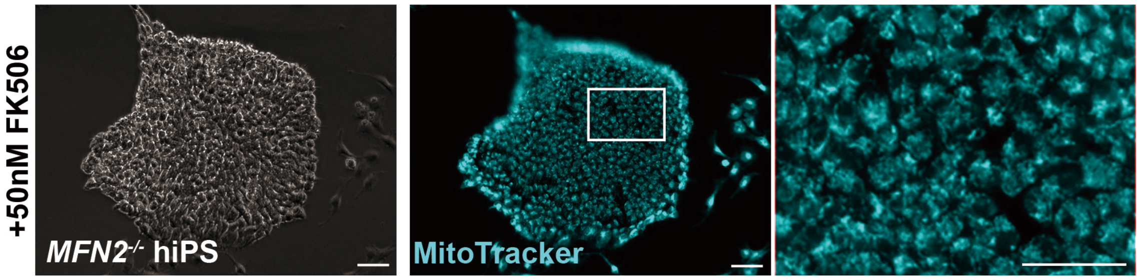


**Supplementary file 10. Mitochondrial morphological changes in MFN2-deficient human iPS cells remained unaltered in the presence of FK506, a calcineurin inhibitor.** MFN2-deficient human iPS cells were treated with FK506, and the mitochondrial morphology was monitored using Mitotracker (blue, right panels). Phase contrast is shown on the left panels. Scale bar; 100 μm.

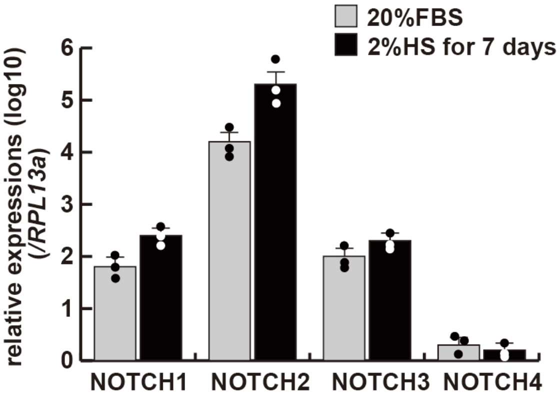


**Supplementary file 11. Notch2 was expressed at the highest level compared to other Notch family members in human myogenic cells.** Relative transcript levels of Notch1~4 genes in growing (20%FBS) and differentiated (2% HS for 7 days) human primary hOM2 cells. FBS; fetal bovine serum, HS; horse serum.

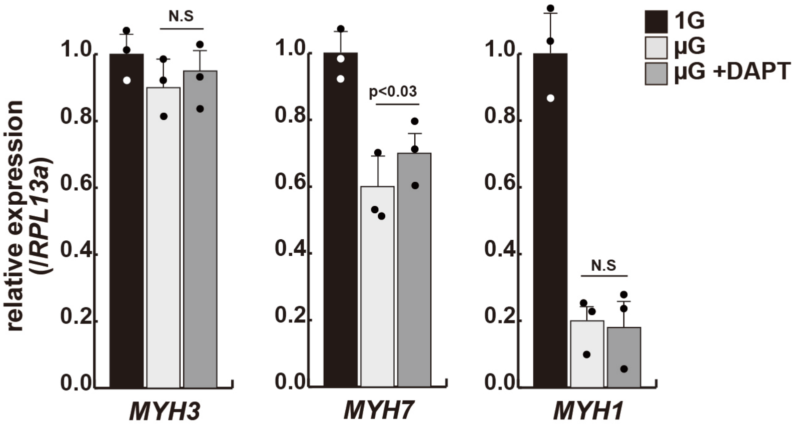


**Supplementary file 12. Myosin heavy chain expressions in differentiated hOM2 myogenic cells with or without DAPT under microgravity.** Relative transcript levels of *MYH3*, *MYH7*, and *MYH1* genes in differentiated human primary hOM2 cells. 1G; under normal gravity, μG; under microgravity, μG+DAPT; with DAPT under microgravity, N.S; not significant.

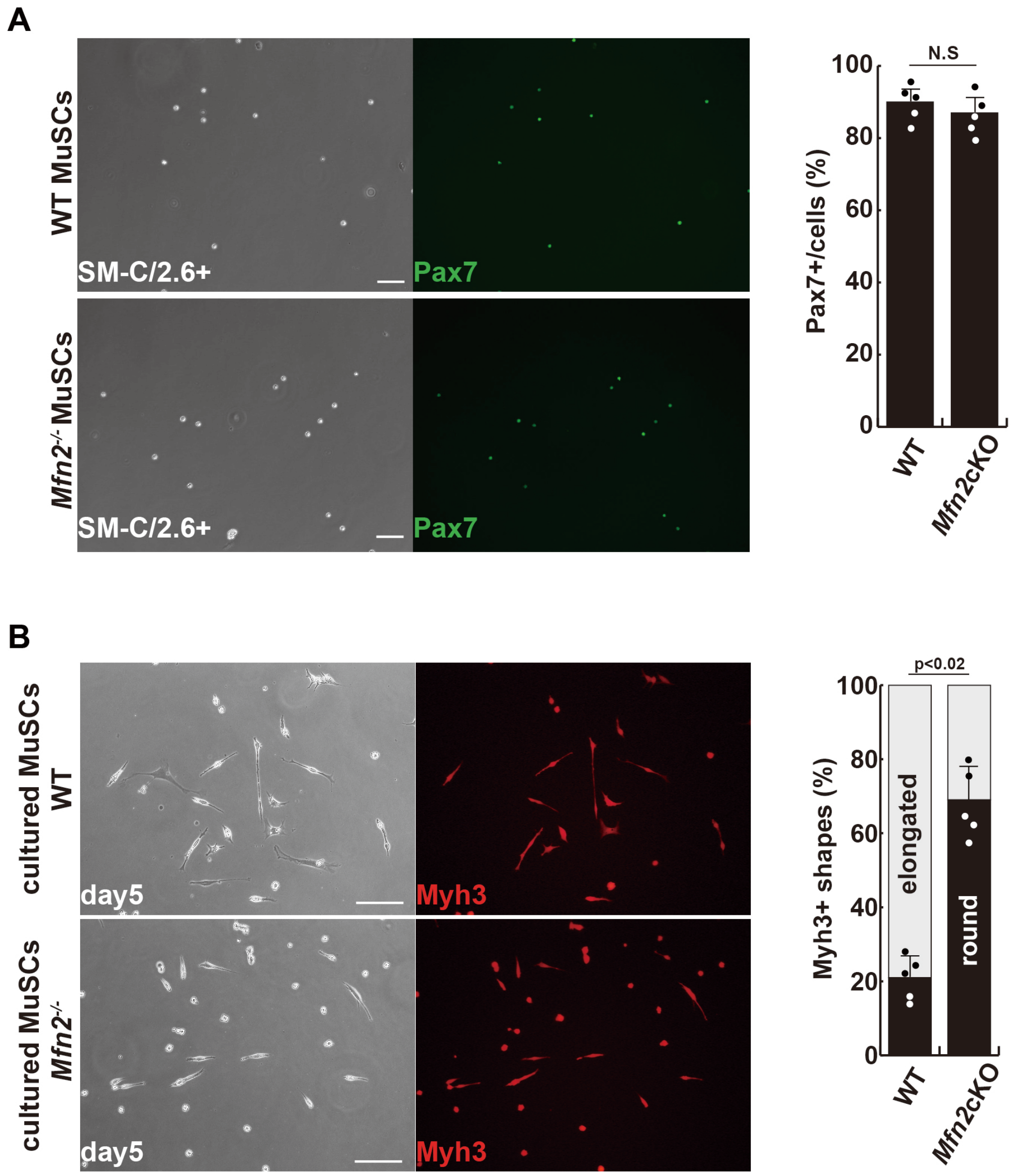


**Supplementary file 13. Isolated muscle stem cells derived from conditional *Mfn2* knockout mice.** (A) Phase contrast (left panels) and Immunofluorescent (right panels) images of isolated muscle stem cells (MuSCs) as SM-C/2.6 positive cells derived from each TA muscle, with anti-Pax7 (Green) as a marker of muscle stem cells, and the proportion of Pax7-positive cells among all isolated cells (right). (B) Phase contrast (left panels) and Immunofluorescent (right panels) images of cultured MuSCs for 5 days with anti-Myh3 (Red) as a marker of differentiated muscle cells (left), and the proportion of Myh3-positive, non-elongated, round-type myogenic cells (right). All error bars indicate ±SEM (n=4). *P*-values are determined by non-parametric Wilcoxon tests for comparisons. N.S; not significant.

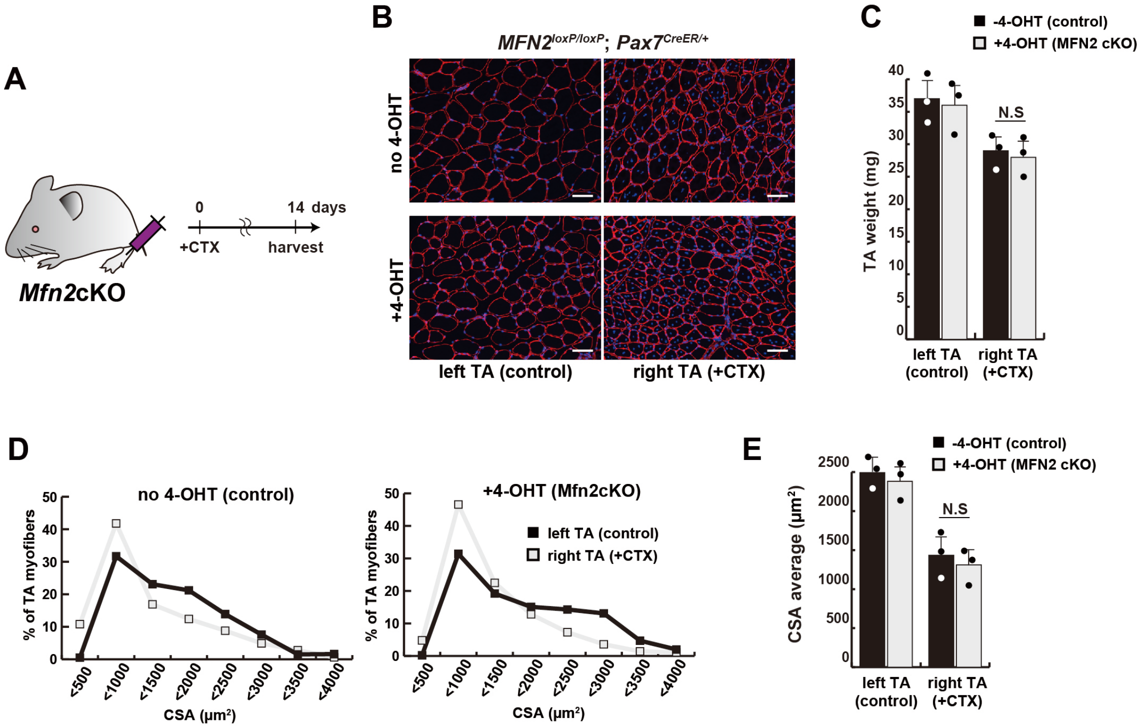


**Supplementary file 14. The regenerative capacity of conditional Mfn2 knockout mice, specifically in muscle stem cells, did not exhibit any significant alteration following a single muscle injury.** (A) Experimental design with Mfn2 conditional knockout mice (Mfn2cKO) for CTX injection and sample collection after 14 days. (B) Immunofluorescence of laminin-2a (red) and DAPI (blue) on transverse sections in the tibialis anterior (TA) muscle of conditional *Mfn2c*KO (*Mfn2*$^{loxP/loxP}$; *Pax7*$^{CreERT2/+}$) mice with or without 4-hydroxytamoxifen (4-OHT) at 12 weeks old of age. (C) TA weight of control left legs and CTX-injected right legs with or without 4-OHT. (D) Distribution of cross-section areas (CSA) of myofibers in each TA muscle. (E) Average CSA of each TA sample. All error bars indicate ±SEM (n=3). N.S; not significant.

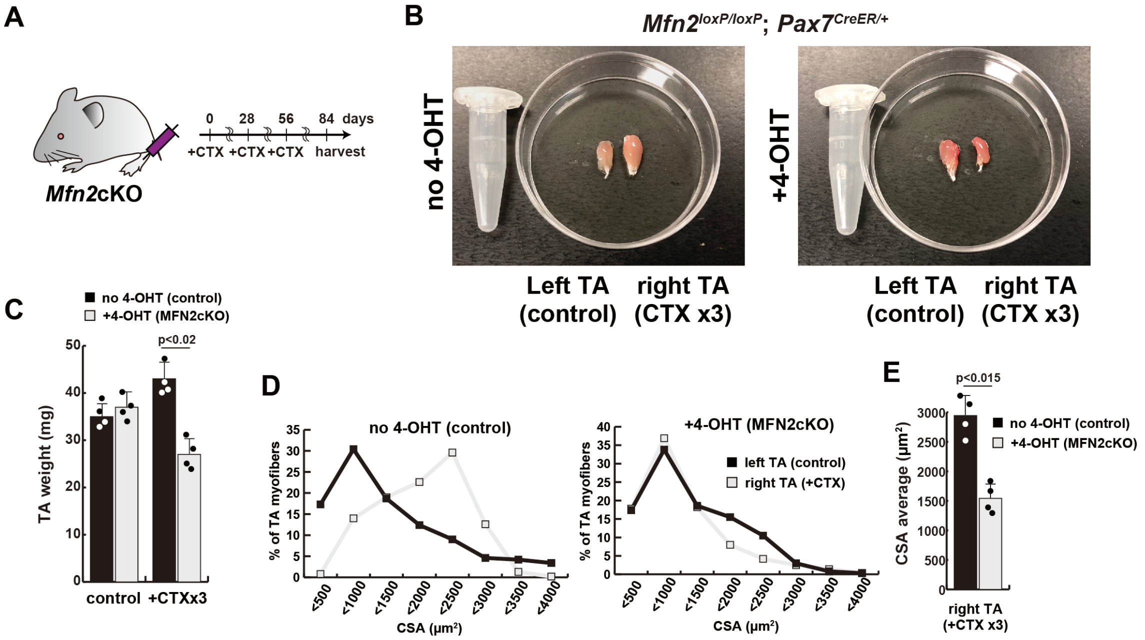


**Supplementary file 15. Reduced muscle hypertrophy in conditional *Mfn2* knockout mice after several CTX-induced muscle injuries.** (A) Experimental design utilizing Mfn2 conditional *Mfn2c*KO (*Mfn2*$^{loxP/loxP}$; *Pax7*$^{CreERT2/+}$) mice, involving 3 rounds of CTX injection to each right TA muscle at 28-day intervals, with sample collection conducted 84 days after the initial CTX injection. (B) Photographs of TA muscles after the dissection. (C) TA weight of control left legs and CTX-injected right legs with or without 4-OHT. (D) Distribution of cross-section areas (CSA) of myofibers in each TA muscle. (E) Average CSA of each TA sample. All error bars indicate ±SEM (n=4). *P*-values are determined by non-parametric Wilcoxon tests for comparisons.

**Supplemenrary file 16**. Primers for the expression analysis by RT-qPCR of the mRNAs indicated.

| Gene | Sequence | Product |
|---|---|---|
| *RPL13A* | 5'-CCCTGGAGGAGAAGAGGAAA-3' | 91bp |
| | 5'-ACGTTCTTCTCGGCCTGTTT-3' | |
| *GAPDH* | 5'-ATGGGGAAGGTGAAGGTCGG-3' | 70bp |
| | 5'-TAAAAGCAGCCCTGGTGACC-3' | |
| *ACTB* | 5'-CACCATTGGCAATGAGCGGTTC-3' | 135bp |
| | 5'-AGGTCTTTGCGGATGTCCACGT-3' | |
| *MFN1* | 5' -GGGTGCTCCTAGGATTATCAGA-3' | 93bp |
| | 5'-TATCTGGCGTTGCTGGAGT-3' | |
| *MFN2* | 5'-TGCCTCAGAGCCCGAGTA-3' | 60bp |
| | 5'-CTGGTACAACGCTCCATGTG-3' | |
| *MYOG* | 5'-GCTCAGCTCCCTCAACCA-3' | 94bp |
| | 5'-ACGTTCTTCTCGGCCTGTTT-3' | |
| *MRF4* | 5'-CAGCTACAGACCCAAACAAGAA-3' | 91bp |
| | 5'-TCCTGGAATGATCGGAAACAC-3' | |
| *MYH1* | 5'-TAAGACCGAGGCAAAAAGGA-3' | 112bp |
| | 5'-TGCATCAGCCAAGCTGTC-3' | |
| *MYH2* | 5'-TTCATGCTGACTGACCGAGA-3' | 197bp |
| | 5'-AGTAGGGGGTTGGCACTGAT-3' | |
| *MYH3* | 5'-GCAGATTGAGCTGGAAAAGG-3' | 167bp |
| | 5'-TCAGCTGCTCGATCTCTTCA-3' | |
| *MYH4* | 5'-GGGGACCCTTGAAGATCAAAT-3' | 150bp |
| | 5'-ATATCTGCAGAAGCCAGTTTGC-3' | |
| *MYH7* | 5'-ACCCTGTTTGCCAACTATGC-3' | 198bp |
| | 5'-CATCACCCCTGGAGACTTTG-3' | |
| *HES1* | 5'-GAAGCACCTCCGGAACCT-3' | 111bp |
| | 5'-GTCACCTCGTTCATGCACTC-3' | |
| *HEY1* | 5'-CATACGGCAGGAGGGAAAG-3' | 125bp |
| | 5'-GCATCTAGTCCTTCAATGATGCT-3' | |
| *HEYL* | 5'-CTTGCAGATGACGGTGGAT-3' | 72bp |
| | 5'-GCTCGGGCATCAAAGAATC-3' | |
| *ID1* | 5'-GTTGGAGCTGAACTCGGAATCC-3' | 145bp |
| | 5'-ACACAAGATGCGATCGTCCGCA-3' | |
| *ID2* | 5'-TTGTCAGCCTGCATCACCAGAG-3' | 150bp |
| | 5'-AGCCACACAGTGCTTTGCTGTC-3' | |
| *ID3* | 5'-CAGCTTAGCCAGGTGGAAATCC-3' | 153bp |
| | 5'-GTCGTTGGAGATGACAAGTTCCG-3' | |
| *ID4* | 5'-GGACCTGTCCAGCCGCGCC-3' | 101bp |
| | 5'-TCAGCGGCACAGAATGCTGTCG-3' | |
| *NOTCH1* | 5'-CGGGGCTAACAAAGATATGC-3' | 88bp |
| | 5'-CACCTTGGCGGTCTCGTA-3' | |
| *NOTCH2* | 5'-TGGTGGCAGAACTGATCAAC-3' | 78bp |
| | 5'-CTGCCCAGTGAAGAGCAGAT-3' | |
| *NOTCH3* | 5'-CCTAGTCCTGGCTCCGAAC-3' | 90bp |
| | 5'-GAGCCGGTTGTCAATCTCC-3' | |
| *NOTCH4* | 5'-ACTGCCTCTGTCCTGATGGA-3' | 105bp |
| | 5'-AACCCACGTCACACACACAT-3' | |
| *MuRF1* | 5'-AAGCCAGTGGTCATCTTGCCGT-3' | 118bp |
| | 5'-CTCCAGACATGGACACTGAGCT-3' | |
| *FBXO32* | 5'-CACTGGTCCAAAGAGTCGGCAA-3' | 158bp |
| | 5'-GCACAAAGGCAGGTCAGTGAAG-3' | |
